# Epigenetic adaptation of beta cells across lifespan and disease: age-related demethylation is advanced in type 2 diabetes

**DOI:** 10.1101/2025.07.02.662814

**Authors:** Elisabetta Manduchi, Hélène C. Descamps, Jinping Liu, Jonathan Schug, Tong Da, Deeksha Lahori, Hilana El-Mekkoussi, Michelle Lee, Eseye Feleke, Diana Bernstein, Chengyang Liu, Ali Naji, Benjamin Glaser, Klaus H. Kaestner, Dana Avrahami

## Abstract

Although the prevalence of type 2 diabetes (T2D) increases with age, most adults maintain normoglycemia despite rising insulin resistance, largely due to the adaptive capacity of pancreatic beta cells to meet increased metabolic demand. However, persistent insulin resistance can lead to beta cell dysfunction and T2D onset. Here, leveraging cell-type-specific methylome data from the Human Pancreas Analysis Program (HPAP), we investigate the epigenomic basis of beta cell adaptation by mapping genome-wide DNA methylation (DNAm) patterns across the human lifespan. In healthy donors, we identify progressive age-related demethylation enriched in *cis*- regulatory elements at beta cell identity and function genes, suggesting that epigenetic remodeling supports functional adaptation to metabolic demand over time. In contrast, alpha cells show the opposite trajectory, with subtle, age-related hypermethylation. In T2D beta but not alpha cells we observed further demethylation compared to healthy controls, underscoring a unique capacity of beta cells to respond to changes in metabolic demand. Together, our findings suggest that DNAm remodeling in healthy beta cells reflects a long-term adaptation to metabolic demand, which in T2D is accelerated as part of a compensatory response that ultimately fails under sustained insulin resistance.

## Introduction

Aging is commonly described as the gradual decline in physiological functions and metabolic regulation, a process observed across the animal kingdom. However, while physiological deterioration predominantly occurs in the later stages of life, recent studies suggest that the aging process can also include phases of physiological adaptation and even functional improvement. During human adulthood, many physiological functions remain stable for decades, reflecting ongoing adaptation. For instance, older adults maintain the ability to improve cardiovascular fitness and cardiac output in response to aerobic exercise^1^, enhance endothelial function to reduce arterial stiffness^2^, and sustain daily energy expenditure^3^.

Pancreatic beta cells play a crucial role in regulating glucose homeostasis by secreting insulin in response to elevated blood glucose levels. Failure to adequately compensate for insulin resistance can lead to persistent hyperglycemia and, eventually, type 2 diabetes (T2D)^4^. Although insulin resistance and elevated blood glucose levels tend to worsen with age^4–6^, most older individuals maintain normoglycemia. This suggests that beta cells, which are long-lived and rarely proliferate in adulthood^7–10^, must sustain function over decades by continually adapting to fluctuating metabolic demands. Indeed, beta cells can adapt during physiological insulin-resistant states, such as pregnancy and puberty, by expanding their mass and enhancing their capacity to secrete insulin^11,12^. Our prior transcriptomic analysis of human beta cell ontogeny supports the notion that beta cells activate genes critical to their function to sustain enhanced activity with age^13^. This is further corroborated by a recent report from Shrestha and colleagues which highlights a beneficial adaptive response in beta cells throughout adulthood, associated with enhanced glucose-stimulated insulin secretion (GSIS), that persists at least until age 60^14^.

DNA methylation (DNAm), a relatively stable epigenetic mark, exhibits cell-specific patterns in regulatory regions, helping to maintain cell-type-specific expression programs across multiple cell generations and throughout life^15^. Despite its stability, DNAm patterns are altered gradually over the human lifespan, with changes at specific sites serving as an epigenetic clock and providing one of the most accurate predictors of chronological age^16^. While numerous studies have shown DNAm alterations in the context of cancer^17^, far fewer have explored how factors related to metabolic diseases impact DNAm and, more specifically, how these methylation changes may contribute to the development of T2D. Previous studies have reported DNAm alterations in the promoters and enhancers of genes associated with insulin secretion in whole islets of deceased organ donors with T2D^18–20^. However, as the composition of human islets varies dramatically both within and between individuals^10^, with alterations in the alpha to beta cell ratio in T2D patients due to reduced beta cell mass^21,22^ or possibly alpha cell expansion^23^, it is essential to analyze T2D methylomes in a cell type-specific manner. This is further underscored by the distinct global methylation patterns between alpha and beta cells^24,25^, and by the unique DNAm alterations observed in beta cells across lifespan and in T2D, as described below.

In this work, we leveraged the islet molecular phenotyping data generated by the Human Pancreas Analysis Program (HPAP; https://hpap.pmacs.upenn.edu)^26,27^. To investigate epigenetic adaptation in beta cells across the human lifespan, we analyzed genome-wide DNAm patterns in sorted beta and alpha cell populations from control (CTL) donors of various ages (here and throughout, ‘control’ refers to non-diabetic, autoantibody-negative individuals). This analysis revealed age-associated demethylation in beta cells, particularly in regulatory regions near islet transcription factor (TF) binding sites. Although these regions exist in a hypomethylated state in beta cells, maintaining that state may require active demethylation, which counteracts maintenance methylation^28,29^. This ongoing process may help sustain gene expression programs essential for beta cell function in the face of ongoing metabolic demand. In contrast, alpha cells exhibited a subtler and opposite age-related trend, with a predominance of hypermethylation even in active regulatory regions.

Interestingly, we noted a beta cell-specific additional demethylation process in T2D as compared to CTL, suggestive of an intensified compensatory response. In contrast, alpha cells from diabetic donors exhibited minimal methylation changes when compared to healthy donors. Our findings suggest that, in healthy individuals, the beta cell epigenome supports lifelong adaptation to metabolic demand. In T2D, however, enhanced demethylation likely reflects mounting stress on beta cells as diabetes progresses, representing an intensified but unsustainable adaptive response.

## Results

### DNA methylation patterns distinguish alpha and beta cell regulatory landscapes

DNAm plays a crucial role in gene silencing, which is essential for maintaining cell type identity and cellular expression programs throughout the lifespan and across cell divisions^15^. This stable methylation pattern is established by regulated processes of methylation and demethylation, and preserved by the activity of maintenance methyltransferase (DNMT1), as well as *de novo* methyltransferases such as DNMT3A^30,31^. To confirm that our dataset captures robust cell identity differences, quality control clustering of all 154 HPAP methylome samples shows clear separation between alpha, beta and exocrine cells regardless of sex, ancestry, age, diabetes status, and assay type, with the outliers in each branch corresponding to samples with a low purity score (Fig.S2).

To characterize the cell type-specific methylation landscapes of pancreatic alpha and beta cells in our cohort, we compared methylomes from 14 CTL donors with high sample purity in both cell types (average purity score of 86% (SD 6%) and 91% (SD 7%) for alpha and beta cells respectively; see Table.S1). The donors in this comparison represented an age range from 26 to 58 years. Principal component analysis (PCA) of methylation levels revealed that the sample’s cell identity accounted for 44% of the variance (Fig.1A). Comparison of methylation profiles between alpha and beta cells identified 78,389 Differentially Methylated Regions (DMRs) with FDR<0.01 and |delta|>10% (Fig.1B), whose delta (average DNA methylation difference) distribution is shown in Fig.1C.

**Figure 1.**
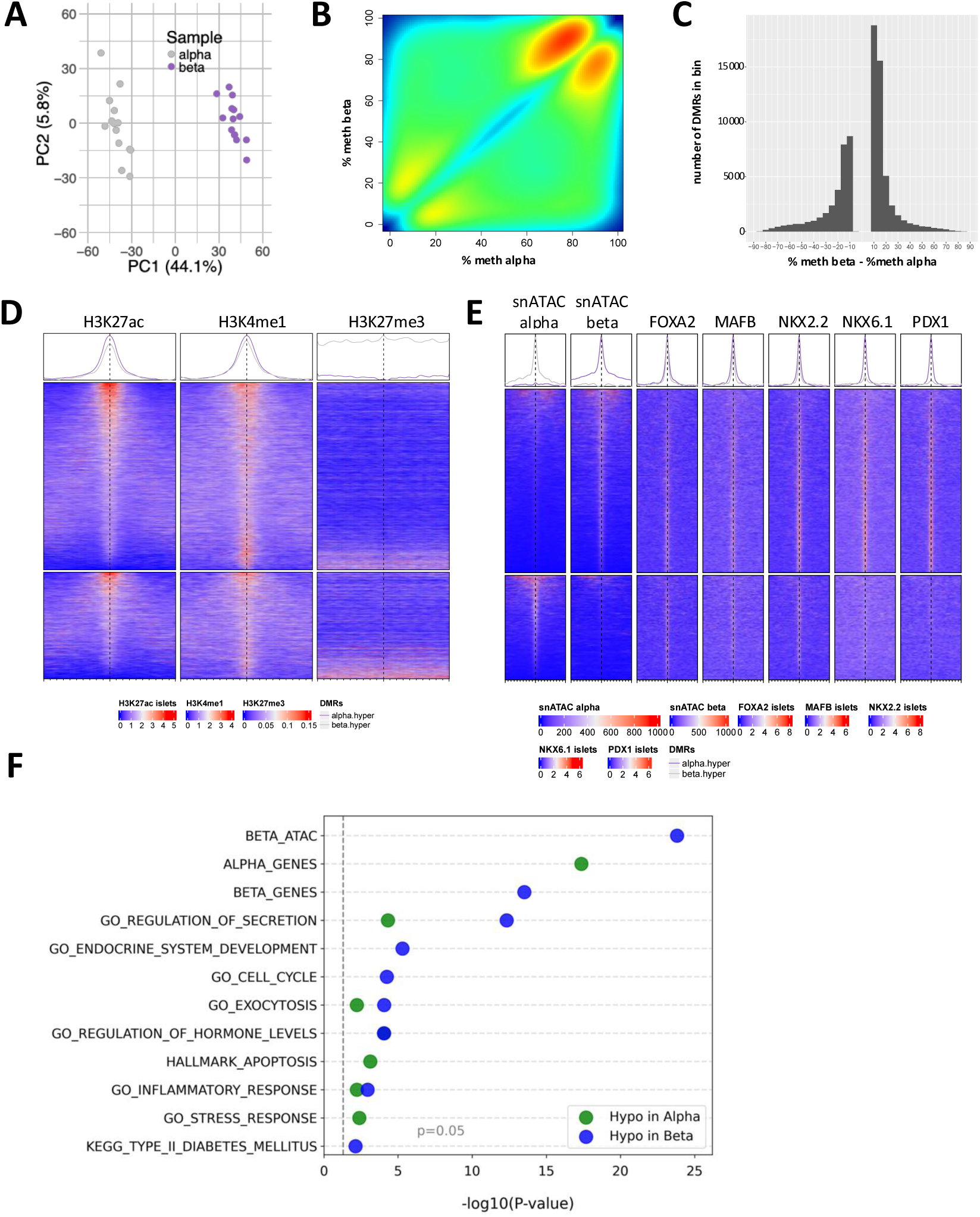
DNA methylation distinguishes alpha and beta cells in association with their cell type-specific regulatory landscapes. (**A**) Scatterplot of the first two principal components from principal component analysis (PCA) of DNAm CTL alpha and beta cells, showing separation by cell type along PC1. (**B**) Smooth scatterplot of beta versus alpha cell methylation values (as %) in CTL donors for 78,389 differentially methylated regions (DMRs) with an FDR<0.01 and |delta|>10%. (**C**) Distribution of the difference in average methylation between alpha and beta cells from the CTL donors DMRs shown in (**B**). X-axis: % difference (beta - alpha); Y-axis; number of DMRs per bin. (**D-E**) Enrichment plot of signal intensity for (**D**) islet histone marks (ChIP-Seq^32,57^ and (**E**) chromatin accessibility (snATAC-Seq, HPAP) and TF binding sites (whole islet, ChIP-Seq^32^) across top alpha-beta DMRs (FDR < 0.01, |delta| > 40%). Each row shows the signal over the 10kb region centered at one of 7,522 DMRs. Top: regions hypomethylated in beta cells; bottom: regions hypomethylated in alpha cells. (**F**) Gene set enrichment for hypomethylated DMRs in alpha (green) and beta (blue) cells, based on DMR-distal genes (FDR < 0.01, |delta| > 40%, CTL alpha vs. beta).

We next focused on the 7,522 DMRs with the strongest effect sizes (|delta|>40%) and analyzed the signals from published islet ChIP-Seq and alpha and beta snATAC-Seq experiments across 10 kb regions centered at these DMRs (see methods). This analysis revealed that regions hypomethylated in both beta and alpha cells were enriched for active chromatin marks and depleted of repressive marks (Fig.1D). Consistently, hypomethylated regions in beta and alpha cells overlapped with open chromatin in each respective cell-type (Fig.1E). Finally, analysis of published TF binding data for whole islets^32^ revealed that binding sites for *FOXA2*, *MAFB*, and *NKX2.2*, which are expressed in both alpha and beta cells, are mostly hypomethylated in both cell types, whereas binding sites for *NKX6*.*1* and *PDX1*, which are specific to beta cells, are predominantly hypomethylated in beta cells (Fig.1E). These findings further demonstrate that alpha and beta cell specific hypomethylated regions are enriched in cell type-specific regulatory elements, extending prior observations^33^.

To test which gene sets are associated with hypomethylation in alpha and beta cells, we applied a hypergeometric test using DMRs selected with thresholds of FDR<0.01 and |delta|>40%, considering genes annotated as DMR-distal, as described in *Methods*. This analysis showed that while common biological pathways important for both cell types, such as regulation of secretion and exocytosis, are associated with hypomethylation in both alpha and beta cells, other gene sets are specifically associated with hypomethylation in either cell-type (Fig.1F). For example, genes critical for beta cell function, such as *ADCYAP1*, *IAPP*, *INS*, *ABCC8*, and *SYT7* (‘BETA_GENES’ in Fig.1F and Table S9), as well as beta cell-specific transcription factors including *PDX1*, *NKX6.1*, *MNX1*, *PAX6*, and *SIX3* (‘GO_ENDOCRINE_SYSTEM_DEVELOPMENT’ in Fig.1F and Table S9) are associated with hypomethylation in beta cells. In contrast, hypomethylation in alpha cells is associated with alpha cell function genes such as *GCG*, *DPP4*, *FAP*, *FEV*, and *ADRB1* (Beta-1 Adrenergic Receptor), the latter known to stimulate glucagon secretion under adrenergic stress^34^ (‘ALPHA_GENES’ in Fig.1F and Table S10).

### Age-progressive DNA methylation loss in beta cell regulatory regions

Beta cells are long-lived cells that must continually adapt to fluctuating metabolic demands^35^. These ongoing adaptations may be supported by the accumulation of epigenetic changes over time. Unlike gene expression, which is highly variable across individuals and becomes increasingly heterogeneous with age^36^, DNAm provides a more stable epigenetic signature that reflects cellular identity and differentiation state. While age-related DNAm changes have been documented in blood and brain tissues^37,38^, they have not been studied in human beta cells. To address this knowledge gap, we analyzed DNAm profiles from pancreatic beta cells collected from 21 HPAP CTL donors across a broad age spectrum from 8 to 64 years old. UXM- estimated purity scores had a mean of 91% and SD of 6% (see Table.S2 for characteristics of the samples used in this comparison). PCA of methylation levels indicated that PC2 was significantly correlated with age (R^2^=0.62, p<0.001) (Fig.2A-B). In this comparison we analyzed age as a numeric covariate, and to measure effect size we defined delta as the difference in average methylation between the group of older donors aged 39-58 and that of younger donors aged 18-35 (see *Methods*). This comparison identified 10,045 DMRs at the thresholds of FDR<0.01, |delta|>5%, with a striking 94% (9,460 DMRs) showing hypomethylation with age (Fig.2C). Although the magnitude of methylation changes was generally modest, with most DMRs showing |delta| values <30% (Fig.2D), the differences were consistent and statistically significant. Thus, despite the small effect size, aging in beta cells is associated with a robust and widespread hypomethylation signature.

**Figure 2.**
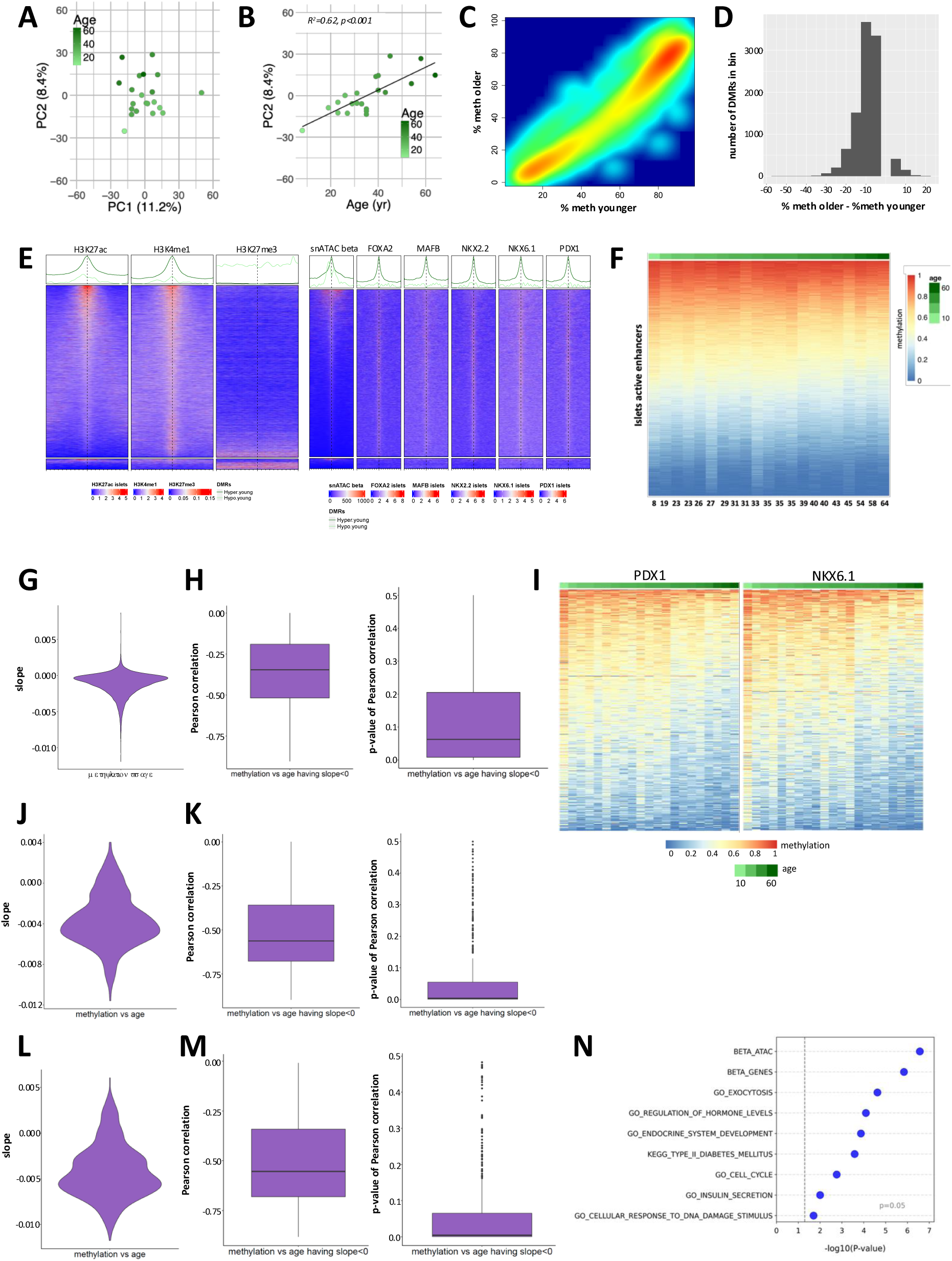
Progressive loss of DNA methylation at regulatory regions in beta cells with age. (**A**) Scatterplot of the first two principal components from PCA of DNAm in CTL beta cells, colored by age (lighter green = younger, darker green=older), showing an association between PC2 and age. (**B**) Scatterplot of PC2 from (A) versus age, illustrating a strong correlation (R^2^=0.62, p<0.001) between the two variables. (**C**) Smooth scatterplot of beta cell methylation values (as %) in older (39-58) vs. younger (18-35) CTL donors across 10,045 age-associated DMRs (FDR<0.01 and |delta| >5%), illustrating age- related demethylation. (**D**) Distribution of average methylation differences between older and younger donors for the DMRs shown in (C). X-axis: % difference (older – younger); Y-axis: number of DMRs per bin. (**E**) Enrichment plot of signals from islet histone marks (left) and chromatin accessibility (snATAC-Seq) and TF binding (ChIP-Seq, whole islets) (right) across 10,045 age-associated DMRs (FDR<0.01 and |delta| >5%) in beta cells from CTL donors. Each row shows the signal over the 10kb region centered at one DMR. Top: regions hypomethylated in older donors; bottom: hypomethylated in younger donors. (**F**) Heatmap of average beta cell methylation at islet active enhancer regions^32^ (rows) across 21 HPAP CTL donors (columns), ordered by age, illustrating progressive demethylation with age. Methylation values are shown as fractions between 0 and 1. (**G**) Violin plot of the slopes of linear fits to methylation versus age, corresponding to the regions in (F). Mean slope=-0.0012, median slope=-0.0008. (**H**) Boxplot of Pearson correlation coefficients (left) and corresponding p-values (right) for regions with negative methylation-age slopes from (F). (**I**) Heatmaps of average beta cell methylation at *PDX1* and *NKX6.1*^32^, showing the 500 peaks with the greatest variability across CTL donors, ordered by age. Note the demethylation trend with age. (**J**) Violin plot of the slopes of linear fits to methylation versus age, corresponding to the *PDX1* regions in (I). (**K**) Boxplot of Pearson correlation coefficients (left) and corresponding p-values (right) for the regions with negative methylation-age slopes from (J). (**L**) Violin plot of the slopes of linear fits to methylation versus age, corresponding to the *NKX6.1* regions in (I). (**M**) Boxplot of Pearson correlation coefficients (left) and corresponding p-values (right) for the regions with negative methylation versus age slopes from (L). (**N**) Gene set enrichment for hypomethylated DMRs in beta cells, based on DMR-distal genes (FDR<0.01 and |delta|>10%) from the CTL beta age comparison.

Next, we analyzed the signals from islet ChIP-Seq and beta snATAC-seq experiments across 10 kb regions centered at these age-related beta cell DMRs (see *Methods*) (Fig.2E). Our analysis revealed that regions hypomethylated with age are predominantly located in beta cell-specific active regulatory chromatin domains, as evidenced by significant enrichment for H3K27ac, H3K4me1, beta-ATAC regions, and binding sites for islet and beta cell-specific TFs such as *NKX6.1* and *PDX1*. To validate these findings, we assessed the average methylation levels within previously identified islet active enhancer regions^32^, which revealed subtle progressive demethylation with age (Fig.2F). The distribution of slopes from linear fits of methylation versus age in these enhancer regions, representing the rate of change of DNAm per year, is clearly shifted towards negative values (Fig.2G). Although the absolute values of the slopes are modest, they reflect a consistent directional trend, with 82% of the 28,813 fits exhibiting negative slopes and relatively strong negative associations, as indicated by the corresponding Pearson correlation coefficients (median=-0.35, Fig.2H left) and p-values (median p = 0.06; Fig.2H right). Notably, enhancer regions can be actively demethylated by TET enzymes, which also help to protect against age-related functional decline in other tissues such as the brain^28,39^.

To explore the mechanisms driving age-associated demethylation in active enhancers, we examined methylation levels at TF binding sites of *NKX6.1* and *PDX1*, which are enriched in beta cell enhancers^40^, focusing on those showing the greatest variation across age. We observed progressive demethylation with increasing donor age at these sites (Fig.2I), and the distribution of slopes from linear fits of methylation versus age was clearly shifted toward negative values, shown for *PDX1* in Fig. 2J and for *NKX6.1* in Fig. 2L. For regions with negative slopes, the median Pearson correlation was –0.56 with a median p-value of 0.004 for *PDX1* (Fig. 2K), and –0.55 with a median p-value of 0.005 for *NKX6.1* (Fig. 2M).

We next asked whether age-related DMRs in beta cells were enriched near genes associated with specific biological pathways or gene sets. Using DMRs filtered by FDR<0.01 and |delta|>10% and considering DMR-distal genes (as defined in *Methods*), we found that hypomethylated regions in older donors were enriched in beta cell-specific regulatory regions, such as beta cell ATAC peaks, and near genes involved in beta cell function, cell cycle regulation, and, of note, DNA damage response (DDR) (Fig.2N, Table S11). GSEA of HPAP scRNA-Seq data from donors in a similar age range revealed up-regulation of the p53 pathway in older individuals, including increased expression of *CDKN2A*, *CDKN1A*, and *CDKN2B* (Fig.S3A, Table S12), supporting progressive activation of stress response and senescence programs in beta cells with age. In line with previous reports^14^, GSEA also showed age-associated down-regulation of the HALLMARK beta cell gene set as a whole (Fig.S3B, Table S13), although individual genes were not significantly downregulated.

### In contrast to beta cells, alpha cells show progressive DNA hypermethylation with age

For alpha cells, we analyzed 14 samples from CTL donors with a mean UXM-estimated purity of 87% and SD of 5% from donors spanning an age range from 18 to 58 years old (Table S3). PCA of methylation levels indicated that PC2 was significantly correlated with age (R^2^=0.77, p<0.001) (Fig.3A-B). We analyzed the CTL alpha cell age comparison in the same manner as described above for beta cells using the same thresholds of FDR<0.01 and |delta|>5%, and identified 5,005 DMRs, of which, surprisingly, 4,800 (96%) were hypermethylated with age, although the magnitude of methylation changes was modest, with most DMRs showing |delta| < 20% (Fig.3C–D). A subset of age-dependent hypermethylation occurred in active regulatory domains, but a higher proportion occurred in repressed regions, not bound by TFs, as illustrated by the enrichment heatmaps of signals from the islet ChIP-Seq and alpha snATAC-Seq experiments over the 10 kb regions centered at these DMRs (Fig.S4; see *Methods* for details).

**Figure 3.**
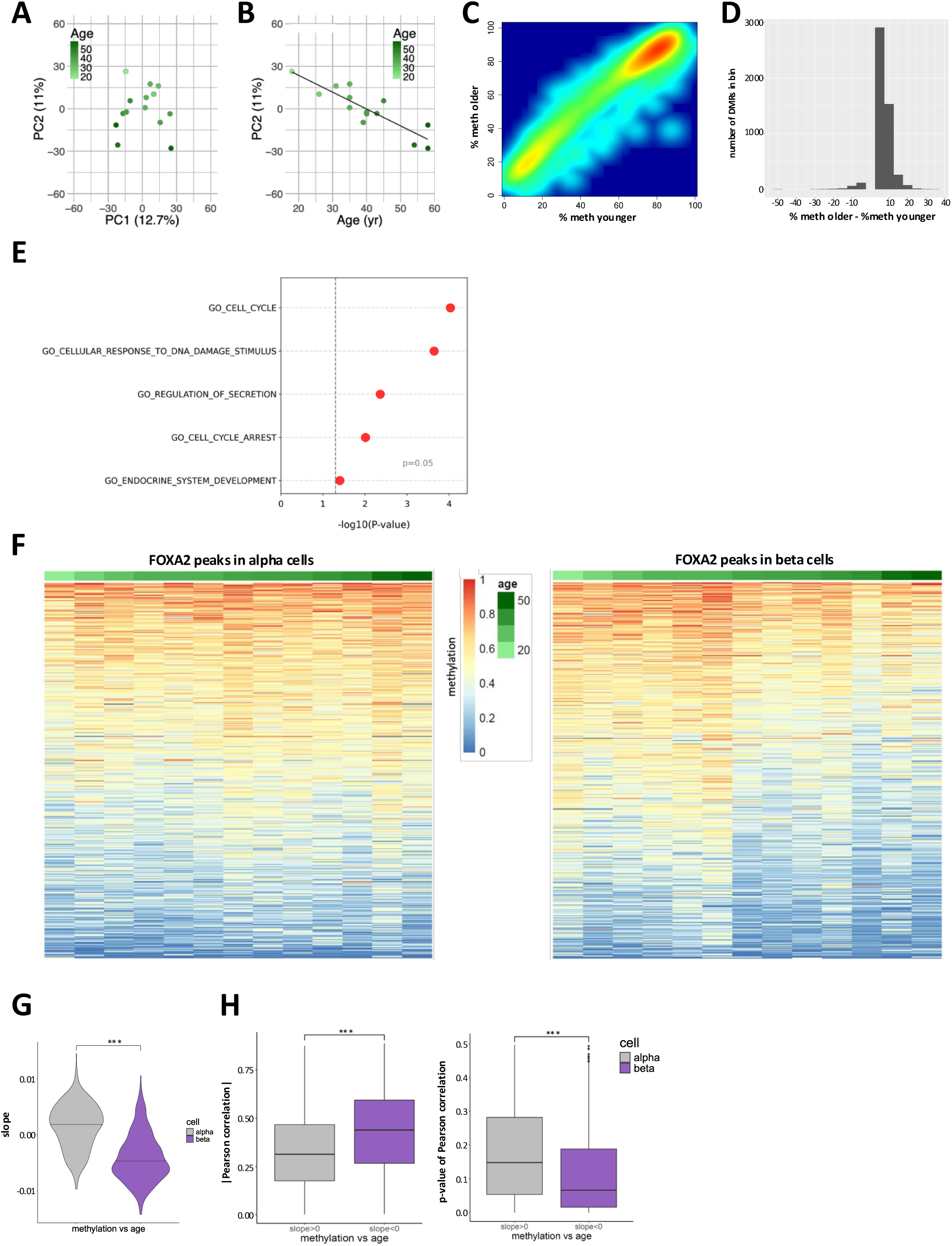
Alpha cells show modest age-related DNA hypermethylation, opposite to the trend in beta cells. (**A**) Scatterplot of the first two principal components from PCA of DNAm in CTL alpha cells, colored by age (lighter green = younger, darker green=older), showing an association between PC2 and age. (**B**) Scatterplot of PC2 from (A) versus age, showing a strong correlation (R^2^=0.77, p<0.001). (**C**) Smooth scatterplot of alpha cell methylation (%) in older (39-58) vs. younger (18-35) CTL donors across 5,005 age-associated DMRs (FDR<0.01, |delta| >5%), illustrating subtle hypermethylation in older donors. (**D**) Distribution of average methylation differences (older – younger) for the DMRs shown in (C). X-axis: % difference; Y-axis: number of DMRs per bin. (**E**) Gene set enrichment for hypermethylated DMRs in alpha cells, based on DMR-distal genes (FDR < 0.01, |delta| > 10%) from the CTL alpha cell age comparison. (**F**) Heatmaps of average CpG methylation at *FOXA2* binding sites^32^ in alpha (left) and beta (right) cells. Shown are the 500 most variable peaks (per cell type) across 13 age-matched donors. (**G**) Violin plots of slopes from linear fits of methylation versus age at the regions shown in (F), for alpha and beta cells. Asterisks indicate Wilcoxon p < 0.001. (**H**) Boxplots of (left) absolute Pearson correlation coefficients and (right) correlation p-values for methylation versus age at regions with slope > 0 in alpha cells and slope < 0 in beta cells from (G). Asterisks indicate Wilcoxon p < 0.001.

We next examined whether age-related hypermethylated DMRs in alpha cells were enriched near specific pathways or gene sets. Using an FDR < 0.01 and |delta| > 10% cutoff, and considering DMR-distal genes, we found that alpha cell-specific hypermethylated regions in older donors were located near genes related to cell cycle regulation and endocrine identity, including transcription factors such as *PAX6* and *NKX2.2* (Fig.3E, Table S14). Analysis of pseudo-bulk alpha cell gene expression data across age did not reveal a correlation with gene silencing or reduced expression with age, possibly due to the low effect size and the higher degree of variability of transcriptomic versus CpG methylation data (data not shown).

To further explore the discrepancy between alpha and beta cells, we compared their methylation levels across ages at binding sites for *FOXA2* and *NKX2.2*, two transcription factors expressed at similar levels in both cell types^32,41–44^. We used the same number of available samples (13) for each cell type (mostly common donors and, when not available, aged-matched donors) and, for each cell type separately, considered the 500 binding sites whose methylation varied most across ages, i.e. focusing on sites most affected by age in that cell type. As illustrated in Fig.3F and Fig.S5, we observed subtle progressing demethylation of these regions with age in beta cells, whereas the trend in alpha cells was opposite. The distribution of slopes from linear fits of methylation versus age in these regions is shifted towards positive values in alpha cells and negative values in beta cells (with the shift more evident at *FOXA2* binding sites than at *NKX2.2* binding sites in alpha cells (Fig.3G and Fig. S6A). Moreover, the regions with negative slopes in beta cells correspond to significantly better linear fits than the regions with positive slopes in alpha cells (Fig.3H and Fig.S6B-C).

Together, these findings demonstrate that despite being closely related cell types residing within the same functional unit, beta and alpha cells undergo opposing age-related DNA methylation trajectories. Beta cells exhibit progressive hypomethylation at regulatory regions, particularly at TF binding sites, potentially reflecting their adaptive, metabolically responsive nature, whereas alpha cells do not show this demethylation process but instead display gradual hypermethylation, predominantly in non-active chromatin regions.

### T2D is associated with premature hypomethylation in beta cells

A recent study using medium-throughput DNAm and RNA-seq analysis of whole human islets reported DNAm changes associated with impaired insulin secretion and reduced expression of beta cell-related genes in T2D donors^20^. However, since islet cell composition varies between individuals^10^, and T2D islets often exhibit altered alpha to beta ratios due to beta cell loss^21,22^ or possibly alpha cell expansion^23^, whole-islet analysis can obscure cell type-specific epigenetic signals. This is especially relevant given the marked differences in DNAm profiles between alpha and beta cells and the opposing age-related methylation changes between the two cell types reported above. To capture beta cell-specific changes in T2D, we analyzed beta cell samples from 12 CTL and seven T2D donors, with similar ages ranging from 33 to 64 years. All samples had high UXM-estimated purity, with a mean of 91% (SD = 7%) for CTL and 89% (SD = 4%) for T2D donors (see Table S4 for characteristics). PCA of methylation levels revealed an association between PC1 and disease status (Wilcoxon p-value=0.04; Fig.4A).

**Figure 4.**
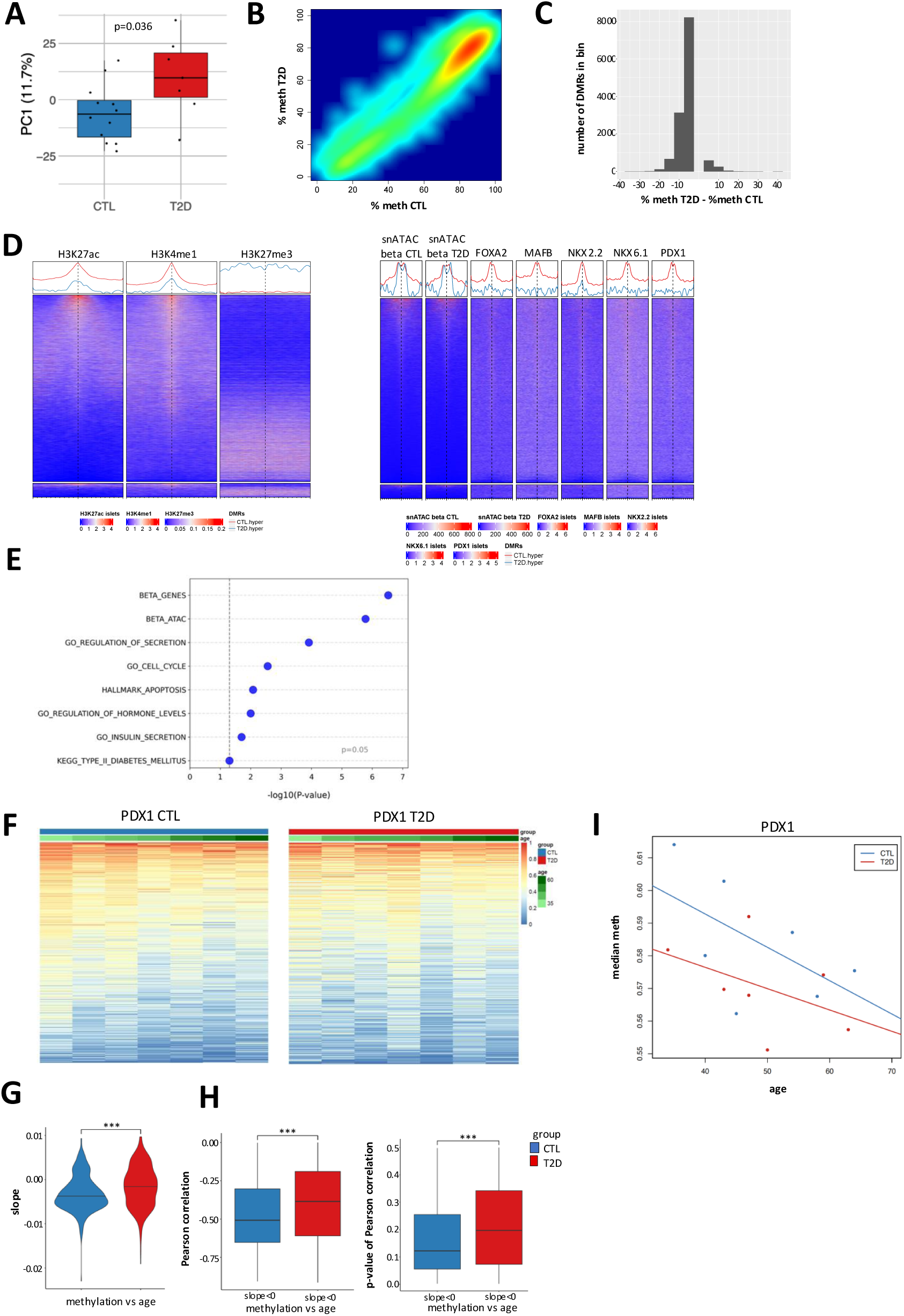
Beta cells exhibit accelerated DNA demethylation in T2D. (**A**) Boxplots of the first principal component from PCA of DNAm in beta cells from CTL and T2D donors. (**B**) Smooth scatterplot of beta cell methylation (%) in T2D versus CTL donors across 13,077 DMRs (FDR < 0.05, |delta| > 5%), illustrating subtle hypomethylation in T2D. (**C**) Distribution of average methylation differences (T2D – CTL) for the DMRs shown in (B). X- axis: % difference; Y-axis: number of DMRs per bin. (**D**) Enrichment plot of signals from islet histone marks (left) and chromatin accessibility (snATAC-Seq, beta cells) and TF binding (ChIP-Seq, whole islets; right) across 13,077 DMRs (FDR<0.05, |delta| >5%) from the T2D vs. CTL beta cell comparison. Each row shows the signal over the 10kb region centered at one DMR. Top: regions hypomethylated in T2D; bottom: hypomethylated in CTL beta cells. (**E**) Gene set enrichment for hypomethylated DMRs based on DMR-distal genes (FDR < 0.05, |delta| > 10%) from the T2D vs. CTL beta cell comparison. (**F**) Heatmaps of average beta cell methylation at *PDX1* binding sites^32^ in CTL (left) and T2D (right) donors. Shown are the 500 most variable peaks (separately, for CTL and T2D) across 7 age-matched donors. (**G**) Violin plots of slopes from linear fits of methylation versus age at the regions shown in (F), for beta cells from CTL and T2D donors. Asterisks indicate Wilcoxon p < 0.001. (**H**) Boxplot of Pearson correlation coefficients (left) and corresponding p-values (right) for methylation versus age at regions with slope < 0 from (G). Asterisks indicate Wilcoxon p < 0.001. (**I**) Median methylation values across the *PDX1* binding sites in the fifth quintile of the median methylation levels for 7 CTL (blue) and 7 T2D (red) age-matched donors.

Using FDR < 0.05 and |delta| > 5% thresholds, we identified 13,077 DMRs, the majority of which (12,170; 93%) were hypomethylated in T2D, with most showing modest differences (|delta| < 20%) (Fig.4B–C). About half of the hypomethylated regions in T2D were enriched for H3K4me1, marking primed enhancers, while a subset also carried H3K27ac, suggesting an active regulatory state. The remaining hypomethylated regions were enriched for the repressive mark H3K27me3 (Fig.4D, left). Some hypomethylated regions also showed enrichment for transcription factor binding sites, particularly those bound by *FOXA2, NKX2.2* and *PDX1* (Fig.4D, right).

We then examined DMR-distal genes to DMRs with FDR < 0.05 and |delta| > 10% and found enrichment in T2D-associated demethylation for genes involved in beta cell–related pathways, mirroring the pattern observed in age-associated hypomethylation (Fig.4E, Table S15). In contrast, no significant pathway enrichment was observed for hypermethylated regions, consistent with a less specific, stochastic process (not shown). To assess the potential functional consequences of these methylation changes, we examined gene expression in beta cells from CTL (n = 13) and T2D (n = 15) donors of similar age using HPAP scRNA-Seq data and found that T2D- associated hypomethylated DMRs did not show a direct correlation with up-regulation of their associated genes (data not shown). However, while core beta cell identity genes such as *PDX1*, *NKX6.1*, *MAFA*, *ABCC8*, and *KCNJ11* maintain stable expression levels in T2D donors (Fig.S7A), we observed reduced expression of broader functional pathways, including oxidative phosphorylation, glycolysis, and protein secretion (2Fig.S7B), supporting intact core identity program, but compromised beta cell function in T2D.

Next, we compared average beta cell methylation levels at *PDX1* binding sites using 7 CTL and 7 T2D donors matched for age, selecting the 500 most variable sites across age within each condition. This analysis revealed that, while demethylation at these sites tends to progress with age in CTL donors as was shown above using a larger number of samples (Fig.2I), this trend is less evident in beta cells from donors with T2D (Fig.4F). The distribution of slopes from linear fits of methylation versus age at these regions was significantly more negative in CTL donors compared with T2D (Fig. 4G). The median Pearson correlation coefficients for regions with negative slopes were also significantly more negative in CTL donors (Fig. 4H, left), and their associated p-values trended lower (Fig.4H, right). While this analysis is based on a relatively small number of age-matched donors and should be interpreted with caution, the consistent differences in slope direction and correlation values across matched CTL and T2D samples support a more robust age-related demethylation process in non-diabetic donors at these regions. The lack of a clear age-related demethylation trajectory in T2D donors likely reflects already reduced methylation levels in beta cells early in life, possibly driven by chronic stress during diabetes progression. To illustrate this observation, we plotted the median methylation levels of the same *PDX1* binding sites in T2D and CTL donors matched for age. As shown in Fig. 4I, focusing on the peaks in the fifth quintile of the median methylation levels (∼55–65%), T2D donors consistently exhibit lower methylation than age-matched controls across the age range. This pattern supports the idea that demethylation at these sites occurs earlier in life in T2D beta cells. Similar trends were observed at *FOXA2* and *NKX2.2* binding sites in the fifth quintile (Fig.S8), further supporting this observation.

Finally, we performed an analysis of DNAm changes in alpha cells from 9 T2D and 13 control donors matched for age (33–63 years) and with high UXM-estimated purity (mean 87%, SD 5% for CTL; mean 86%, SD 8% for T2D; Table S5). Despite the larger number of alpha cell samples compared to the beta cell analysis, we identified significantly fewer DMRs (2,281) between T2D and CTL alpha cells using the same thresholds (FDR < 0.05 and |delta| > 5%) (Fig.5A). These DMRs showed a roughly equal distribution of hypo- and hypermethylation (Fig.5B). Most were marked by the repressive histone modification H3K27me3 (Fig5C), while hypermethylated regions in T2D were also somewhat enriched for *NKX2.2* binding (Fig.5D). Neither hypo- nor hypermethylated regions were associated with specific biological pathways (data not shown).

**Figure 5.**
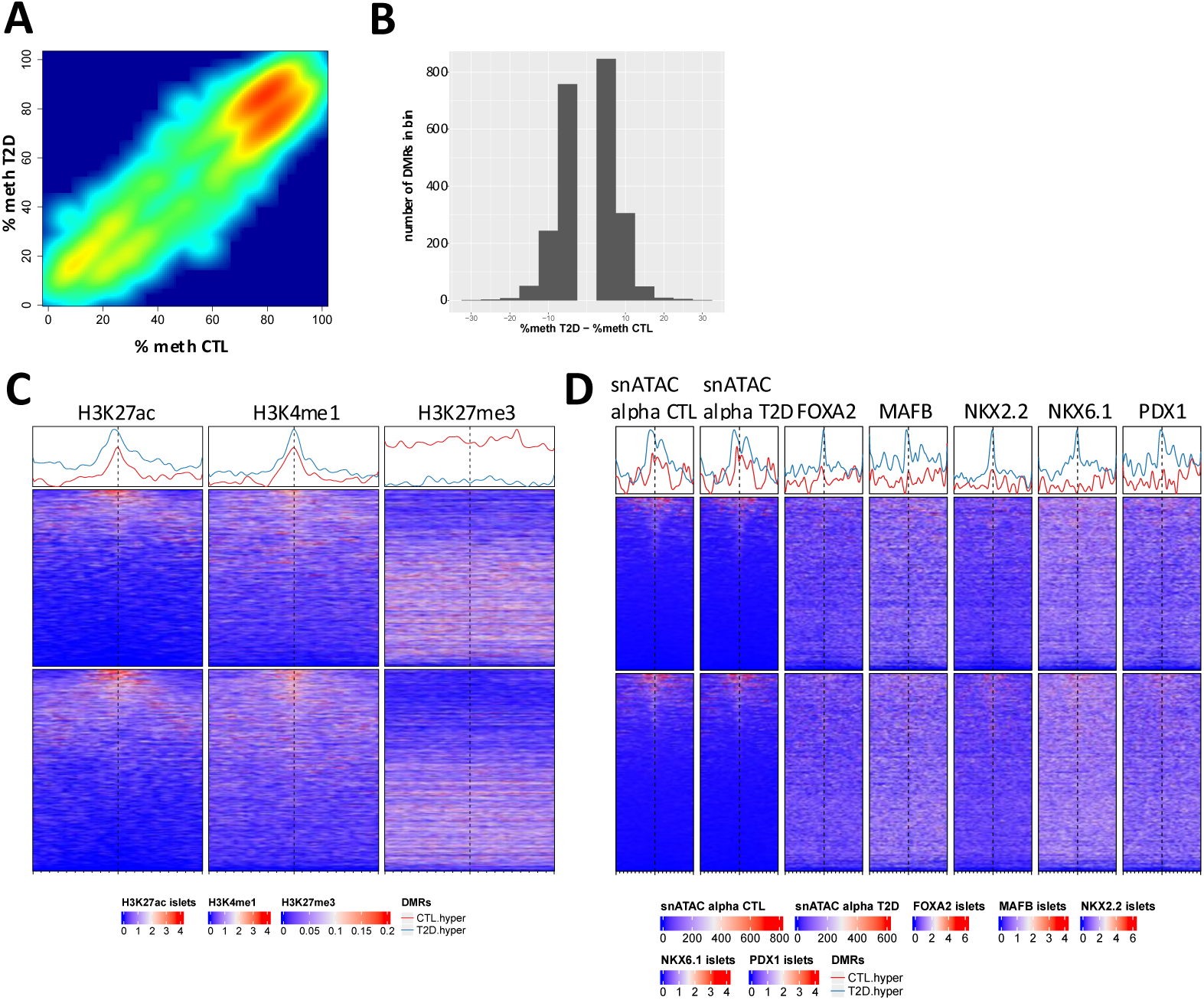
Alpha cells show minimal DNA methylation differences in T2D. (**A**) Smooth scatterplot of alpha cell methylation (%) in T2D vs. CTL donors across 2,281 DMRs (FDR < 0.05, |delta| > 5%), illustrating subtle, bidirectional methylation changes in T2D. (**B**) Distribution of average methylation differences (T2D – CTL) for the DMRs shown in (A). X- axis: % difference; Y-axis: number of DMRs per bin. (**C-D**) Enrichment plots of signals from islet histone marks (C), chromatin accessibility (snATAC- seq, beta cells), and TF binding (ChIP-seq, whole islets) (D) across 2,281 DMRs (FDR < 0.05, |delta| > 5%) from the T2D vs. CTL alpha cell comparison. Each row shows the signal over the 10kb region centered at one DMR. Top: regions hypomethylated in T2D; bottom: hypomethylated in CTL alpha cells.

Altogether, our data demonstrate that beta cells in T2D exhibit an accelerated demethylation pattern resembling age-related changes, predominantly in TF binding sites, a phenomenon not observed in alpha cells, and whose functional consequences remain to be investigated.

## Discussion

This study leverages comprehensive human islet cell methylome datasets generated by the Human Pancreas Analysis Program (HPAP), a unique resource that provides high-resolution, cell type-specific epigenomic profiles from human donors across a range of ages and diabetes status^26,27^. While previous studies have examined DNAm in whole islets, this dataset enables analysis of epigenetic changes in purified alpha and beta cell populations.

Using a subset of HPAP donors with alpha and beta cell samples of the highest purity, we traced DNAm changes associated with age and T2D. Our analysis revealed that beta but not alpha cells from healthy donors undergo progressive demethylation with age at active regulatory regions, particularly at TF-bound *cis* regulatory elements. Due to the limited age range and smaller number of T2D donors, we could not perform an age-related DNAm analysis in T2D. However, a similar demethylation pattern was also noted in T2D as compared to CTL donors, suggesting that T2D beta cells may acquire a hypomethylated state earlier in life, consistent with a previous study showing that premature DNA demethylation in peripheral blood leukocytes is associated with increased risk of T2D^45^. Although the changes are modest in magnitude, they are statistically significant and consistently enriched in functionally relevant regions. The enrichment of hypomethylated regions in regulatory elements suggests a regulated and potentially adaptive process, rather than stochastic drift, targeting loci essential for beta cell identity and function.

DNAm patterns in differentiated cells are generally stable and maintained by the coordinated activity of the maintenance methyltransferase DNMT1 and *de novo* methyltransferases such as DNMT3A to preserve gene silencing^30,31^. However, these patterns can be actively remodeled by TET enzymes, which oxidize 5-methylcytosine (5mC) to 5- hydroxymethylcytosine (5hmC). This intermediate can either initiate active or passive demethylation or persist as a stable epigenetic mark, especially in the context of aging and disease^46^. Importantly, TET-mediated demethylation accumulates at active regulatory regions and TF binding sites^28,47^, where it may counteract DNMT activity and protect enhancers from repressive methylation^28,29,48^. In beta cells, such mechanisms could help sustain enhancer activity and support functionality throughout life. Alternatively, or in addition, the observed demethylation may result from rare replication events triggered by increased metabolic demand throughout life and under diabetic conditions, during which 5hmC-modified sites are not recognized by DNMT1, leading to a loss of CpG methylation in the daughter cells. However, alpha cells, which have been reported to proliferate at higher rate in T2D^23^, do not show substantial hypomethylation. This suggests that replication-dependent passive demethylation alone is insufficient to explain the beta cell-specific pattern, or that beta cells may accumulate higher levels of stable 5hmC over time. Clarifying the relative contributions of these mechanisms will require profiling steady-state 5hmC levels in alpha and beta cells from non-diabetic and diabetic donors.

Beta cell gene expression is influenced by conditions such as ER stress, oxidative stress, and metabolic workload, and gene activation and repression are often reversible, supporting the adaptive capacity of beta cells until chronic stress overwhelms this capacity, leading to failure and progression to T2D^49^. In contrast, DNAm, being a relatively stable epigenetic mark, may not respond to transient metabolic cues or drive rapid changes in gene expression. Indeed, in our study, most age- and disease-associated demethylated regions did not strongly correlate with increased expression of nearby genes. Nonetheless, the specific demethylation process observed in beta, but not alpha, cells, and under diabetic conditions, supports the idea that this represents a cell type specific epigenetic adaptation. In beta cells, demethylation may serve to preserve enhancer activity and sustain cellular responsiveness over time, rather than directly modulate gene expression acutely. Although the functional significance of this demethylation remains to be fully elucidated, its cell-type specificity highlights a unique epigenetic adaptive response in this long-lived, metabolically responsive cell type. However, in the context of T2D, this mechanism may ultimately fail, leading to impaired function despite preserved expression of core identity genes. Future studies should explore whether targeted modulation of the DNAm machinery, particularly TET activity, can enhance beta cell resilience during aging and diabetes.

## METHODS

### Human pancreatic islets

Pancreatic islets were procured by the HPAP consortium under the Human Islet Research Network (https://hirnetwork.org/) with approval from the University of Florida Institutional Review Board (IRB # 201600029) and the United Network for Organ Sharing (UNOS). A legal representative for each donor provided informed consent prior to organ retrieval. The protocols utilized for the human islet sorting, Whole Genome Bisulfite Sequencing (WGBS), single-cell RNA-Seq (scRNA-Seq), and single-nucleus ATAC-seq (snATAC-Seq) assays utilized in this work are available on the PancDB website at https://hpap.pmacs.upenn.edu/explore/workflow/islet-molecular-phenotyping-studies (see Kaestner lab protocols). In addition to libraries generated with WGBS, the methylome data used in this work also comprises libraries generated using EM-Seq following these steps: 100ng DNA input was sheared to 300bp and cleaned with Zymo research kit (#D4014). Quality control was performed on a BioAnalyzer (Agilent) and EM-Seq libraries were generated (New England Biolabs, Ipswich, MA, USA – Catalog number: E7120/E7140). All kits were used following the manufacturer’s instructions.

### Data analysis

#### Methylome

All available (154) HPAP methylome samples, regardless of cell type or donor disease state, were pre-processed together (see Fig.S1 for workflow outline). Adapters were trimmed from paired end reads with cutadapt v4.4^50^. Trimmed reads were aligned to the human hg38, Lambda, and pUC19 genomes using bwa-meth v0.2.7^51^. The resulting sam files were converted to bam files using samtools v1.14^52^, then duplicated reads were marked by sambamba v0.8.1^53^ and resulting bam files sorted and indexed with samtools. We utilized wgbstools v0.2.0^24^ to generate preliminary pat, beta, and tab files from the hg38 alignments while removing reads with low mapping quality, duplicated, or not mapped in a proper pair (using -F 1796 -q 10). We then leveraged the HPAP whole genome sequencing data as input for custom scripts to identify the C>T and G>A variants at CpG positions for each donor and subtracted them from the tab files, also employing bedtools v2.26.0^54^. Utilizing wgbstools again, we generated beta files from the filtered tab files, then computed average coverage, generated bigwig tracks, and segmented the genome using these filtered beta files. In general, the segmentation from wgbstools identifies non-overlapping continuous regions of highly correlated CpG sites similarly methylated in each sample but possibly covarying across conditions. We derived the segmentation from all samples with an average coverage above the median (22X) and required a minimum of 4 CpGs per region and a max region size of 2,000 bp. This segmentation (2,061,859 regions with median size=607 bp and median number of CpGs=8) was then applied to all samples and used in subsequent analyses.

For each sample, we estimated purity utilizing the UXM fragment-level deconvolution algorithm^24^ with reference Atlas.U25.l4.hg38.full.tsv (from https://github.com/nloyfer/UXM_deconv) restricted to the markers for pancreatic cell types, endothelial cells, and monocytes and macrophages. Before applying UXM, we refined the atlas for each donor by filtering out CpG positions where that donor had C>T or G>A variants.

We leveraged methylSig v1.11.0^55^ for differential methylation analyses. We performed several comparisons including (i) alpha versus beta cells in CTL donors, (ii) age (as a numeric covariate) separately in alpha and (iii) in beta cells of CTL donors, (iv) T2D versus CTL donors separately for alpha and (v) beta cells. For each comparison, we selected relevant samples with sufficiently high average coverage and estimated purity. For comparisons between T2D and CTL donors, we selected donors in similar age ranges in the two groups. The characteristics of the samples used in each comparison are summarized in supplementary Tables S1-S5. For each comparison, we removed CpGs covered by less than 10 reads in any of the samples and then aggregated methylation fractions by segmentation regions and removed any region covered by less than 10 reads in any of the samples. We focused on regions not on chromosomes X and Y. To find DMRs, we employed the general model test with DSS and design dependent on the comparison of interest. In comparisons (i) we paired by donor, in comparisons (iv) and (v) we included ancestry in the model. For each comparison, DMRs for further examination were defined based on thresholds for the false discovery rate and the effect size, where the effect size here refers to the absolute value of delta, the difference of average methylation levels. In the case of the age comparisons (where age was treated as numeric covariate), delta refers to the difference in average methylation between the group of donors aged 39-58 and that of donors aged 18-35. The thresholds chosen varied with the comparisons (Table S6) to yield a feasible number of DMRs for follow-up. To functionally annotate DMRs, we defined the following sets of features based on the Gencode Human v43 transcript models:

● 1,500 bp upstream of the Transcription Start Site (TSS)
● 1,501-5,000 bp upstream of TSS
● 5,001-50,000 bp upstream of TSS
● 1,000 bp downstream of TSS
● 1,001-50,000 bp downstream of TSS and downstream of the Transcription End Site (TES). We employed bedtools v2.26.0^54^ to mark the features from each of the above sets overlapping with each DMR.

To explore enrichment of signals from alpha, beta, and islet specific snATAC-Seq and ChIP-Seq experiments, we leveraged EnrichedHeatmap v1.32.0^56^. We downloaded the bigwig files with accessions listed in Table S7 from the Common Metabolic Disease Genome Atlas (https://cmdga.org/). For H3K27me3, we used data from Bramswig and colleagues^57^ lifted over to hg38 with LiftOver^58^. In addition, we generated bigwig files for chromatin accessibility in alpha and beta cells from HPAP snATAC-Seq experiments in CTL and T2D donors as described below. Our plots also leveraged pheatmap v1.0.12 (https://github.com/raivokolde/pheatmap) and ggplot2 v3.5.1^59^.

#### scRNA-Seq

For each donor islet sample, we obtained counts with CellRanger v7.1.0^60^ and utilized Seurat v4.9.9.9041^61^ (complemented by SoupX^62^ and scDblFinder^63^) for clean-up, normalization, pre-processing and data integration. Cells were annotated with two approaches: scSorter^64^ and manual cluster annotation based on selected pancreatic cell markers. Only cells for which the two approaches agreed were assigned to a final cell type.

For each cell type, we utilized all available relevant donors for a comparison of interest, with at least 100 cells of that type and processed with SC3Pv3 chemistry. We generated pseudo- bulk counts and used DESeq2 for differential expression analyses^65^. For the age comparison, we did not treat age as a numeric covariate as we did for DMRs, but we compared age group categories (18-35 vs 39-58 years old) based on DESeq2 recommended guidelines.

#### snATAC-Seq profiles in alpha and beta cells

Chromatin accessibility profiles for alpha and beta cells were generated using HPAP snATAC-Seq datasets. Each dataset was first analyzed using Signac (v1.7.0)^66^ and Seurat (v4.4.0)^61^. A standard processing pipeline was applied (https://stuartlab.org/signac/articles/pbmc_vignette). Nuclei with number of reads in peaks <1,000 or >100,000, TSS enrichment score < 1, or nucleosome signal > 2 were filtered out. Each nucleus was annotated by mapping to an internal multiome reference dataset and nuclei with a mapping score > 0.8 were kept.

Then, to generate cell type specific tracks in each disease group, we used samples from CTL donors with age ≥ 30 and samples from T2D donors. With cell types annotated from the previous steps, fragments associated with cells from each donor and each cell type (alpha and beta) were merged to generate pseudo-bulk tracks. Pseudo-bulk tracks generated with less than 30 cells were removed. A bed file including the start, end, and chromosome information of each read was created for each donor and each cell type. Bam files were constructed using bedToBam v2.30.0^54^ with genome size file provided as part of the GRCh38 reference – 2020-A-2.0.0 available on the 10x Genomics website (https://cf.10xgenomics.com/supp/cell-arc/refdata-cellranger-arc-GRCh38-2020-A-2.0.0.tar.gz). Bam files were indexed using samtools v1.11 index^52^. BAMscale v0.0.5^67^ was used to generate scaled coverage tracks, with parameter -t 4. Lastly, to generate cell type specific tracks per disease group, cell type specific tracks were merged across donors of the same condition using bigWigMerge v2 and the output was sorted and converted to a bigwig file using bedGraphToBigWig v2.8^68^.

#### Gene set analyses

We explored the enrichment of a gene set collection comprising gene sets obtained from MSigDB^69^ and custom gene sets defined in^13^. The gene sets in this collection are provided in Table S8.

To analyze enrichment of gene sets associated with DNAm changes, we employed a hypergeometric test implemented in the Genomica software^70^, considering gene sets with a p-value <0.01 and an FDR <0.05 to be significant. For association of genes to DMRs, we utilized protein coding genes in the unions of the following DMR annotation files defined above: 1,501-50,000 bp upstream of TSS or 1,001-50,000 bp downstream of TSS and downstream of TES. We refer to these genes as ‘DMR-distal’.

To analyze enrichment of gene sets associated with mRNA expression, we used Gene Set Enrichment Analysis (GSEA)^69,71^ on the DESeq2 normalized scRNA-Seq pseudo-bulk counts. The gene sets input in these analyses were the Hallmark gene sets from MSigDB.

## Reporting Summary

Further information on research design is available in the Nature Portfolio Reporting Summary linked to this article.

## Data Availability

Raw data and metadata related to this work are available on the PancDB website https://hpap.pmacs.upenn.edu and on dbGaP (https://www.ncbi.nlm.nih.gov/gap/) with accession phs002465.

## Code Availability

The study was conducted using only publicly available software as outlined above; no custom code was used.

## Acknowledgments

We thank the families of the organ donors and the entire HPAP team for making this study possible. This manuscript used data acquired from the Human Pancreas Analysis Program (HPAP-RRID:SCR_016202) Database (https://hpap.pmacs.upenn.edu/), a Human Islet Research Network (RRID:SCR_014393) consortium supported by NIDDK grants UC4-DK-112217, U01-DK- 123594, UC4-DK-112232, and U01-DK-123716. Further support was provided by NIDDK grant U01-DK-134995 to D.A., B.G. and K.H.K. B.G. and D.A. are supported by grants from the Israel Science Foundation and JDRF (1782/18 and 2982/20), the US-Israel Binational Science Foundation (2019314) and the EU’s Horizon 2020 research and innovation program No 874710. D.A is supported by the Israel Science Foundation (2166/24).

## Author contributions

D.A. and K.H.K conceived and supervised this work. B.G. provided conceptual input. E.M., D.A. and K.H.K. wrote the manuscript. E.M. pre-processed the data, performed most computational analyses, and uploaded the data to PancDB and dbGaP. H.C.D. performed human islet sorting, prepared and sequenced the EM-Seq, scRNA-Seq and snATAC-Seq libraries, performed the PC analyses, and contributed to data interpretation and manuscript outline. D.A. performed gene set analyses and GSEA. J.L. prepared the WGBS libraries. J.S. oversaw the sequencing, provided feedback, and helped with data uploads to PancDB and dbGaP. T.D., D.L., D.B. and H.E.M. contributed to human islet sorting and generated scRNA-Seq and snATAC-Seq libraries. M.L. performed the snATAC-Seq analyses. C.L. and A.N. performed islet isolation from donor tissues. E.F performed additional biological analyses. All authors reviewed and approved the manuscript.

## SUPPLEMENTAL FIGURE LEGENDS

**Figure S1.**
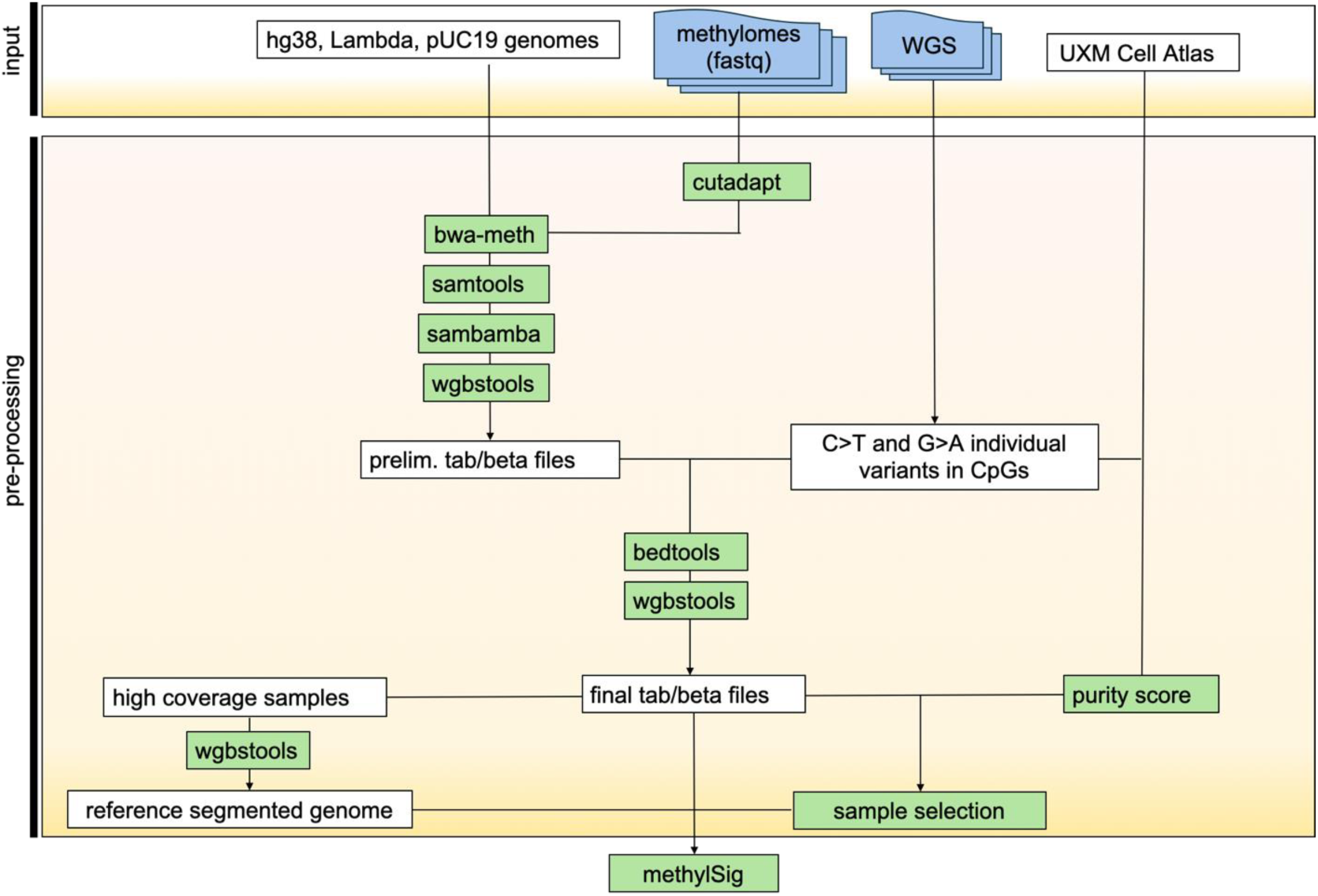
Workflow of the HPAP methylation data pre-processing.

**Figure S2.**
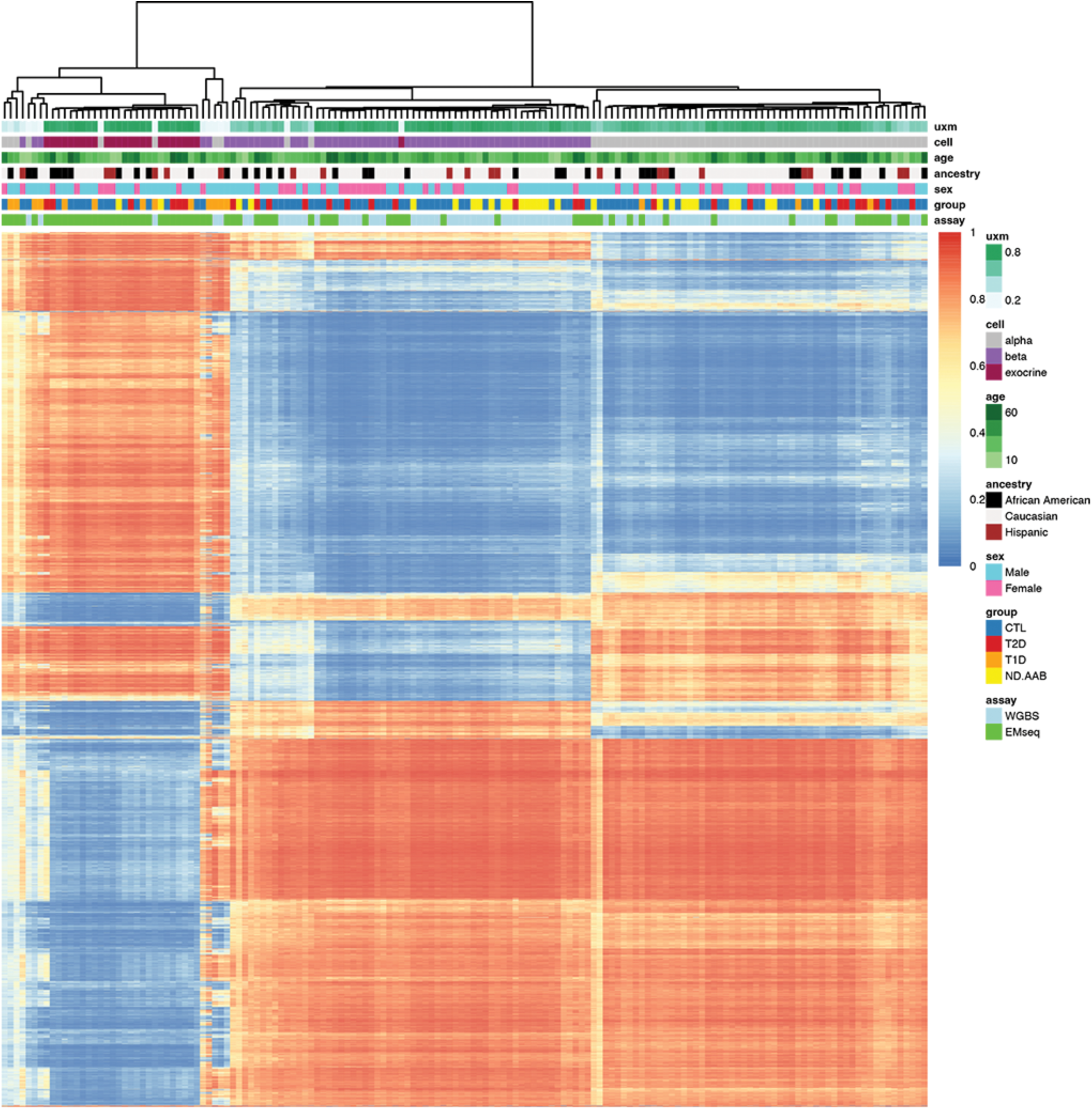
Quality control clustering of all 154 HPAP methylome samples. Rows represent the top 1% most variable segmentation regions (18,972) on chromosomes other than X and Y. Columns correspond to samples. Rows and columns are clustered using Manhattan distance as similarity metrics. Horizontal bars at the top indicate values of several covariates for each sample. ‘uxm’=UXM estimated purity score, ND.AAB’= non-diabetic auto-antibody positive.

**Figure S3.**
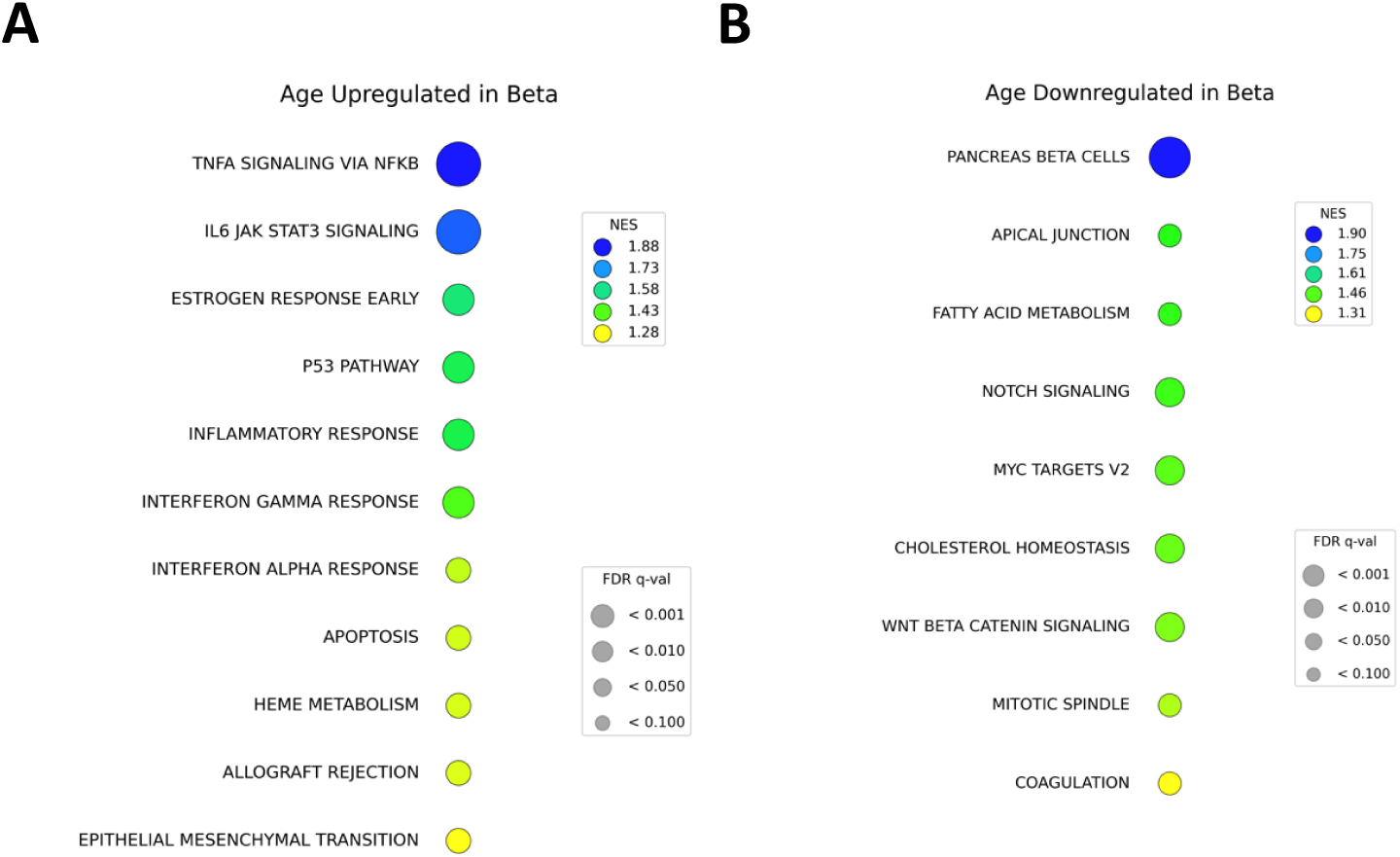
MSigDB Hallmark gene sets identified by GSEA as (**A**) up-regulated with age and (**B**) down-regulated with age in beta cells from CTL donors.

**Figure S4.**
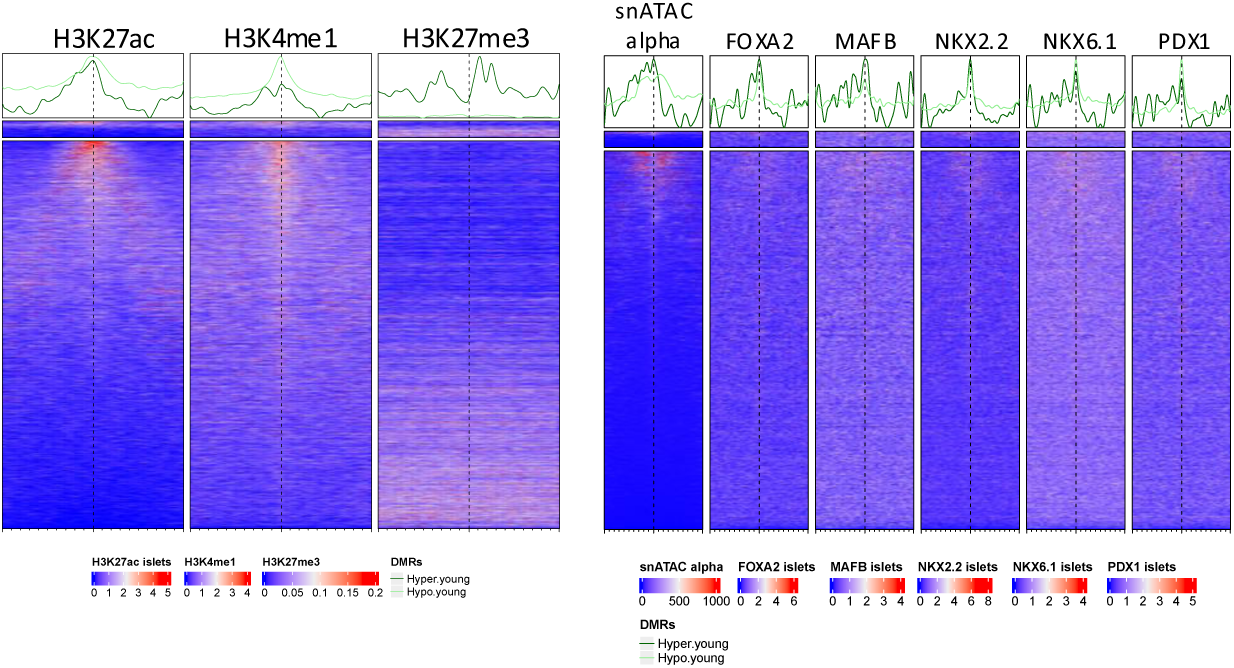
Enrichment plot of signals from islet histone modifications ChIP-Seq (left) and from snATAC-Seq in alpha cells and islet TF ChIP-Seq experiments (right) for the 5,005 age-associated DMRs with FDR<0.01 and |delta| >5% in alpha cells from CTL donors. Each row shows the signal plotted over the 10 kb region centered at one DMR. Top: regions hypermethylated in younger alpha cells; bottom: regions hypermethylated in older alpha cells.

**Figure S5.**
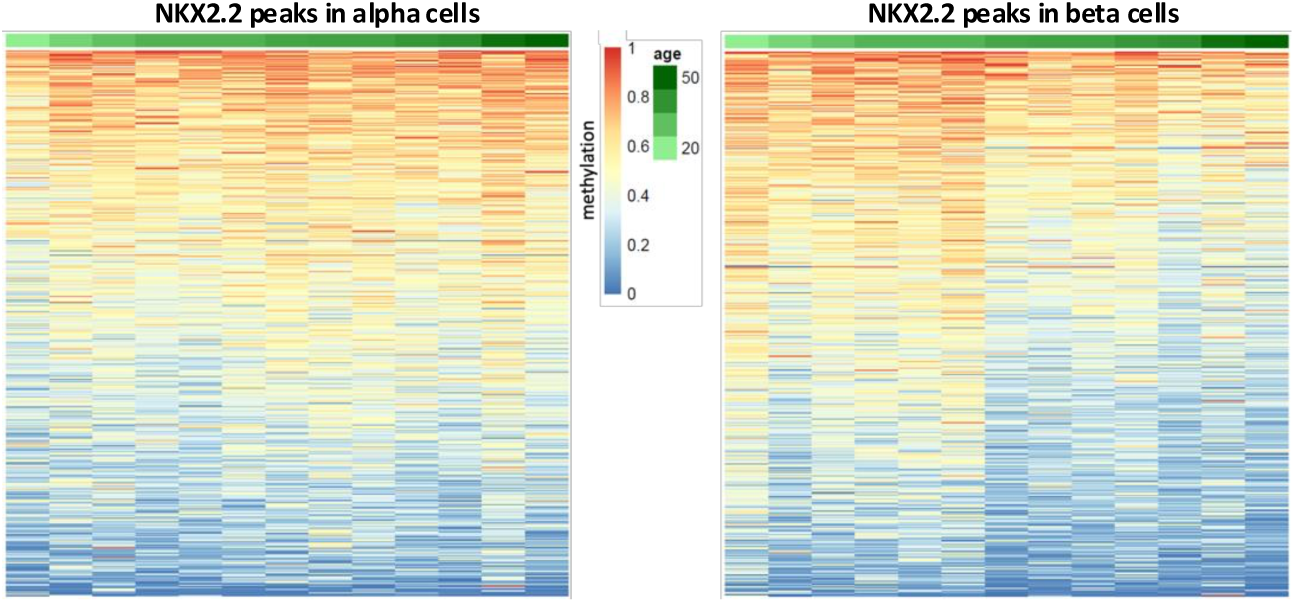
Heatmaps of average methylation for *NKX2.2* peaks^32^ for alpha (left panel) and beta (right panel) cells. Shown are the 500 most variable peaks (separately, for each cell type) across 13 age-matched donors.

**Figure S6.**
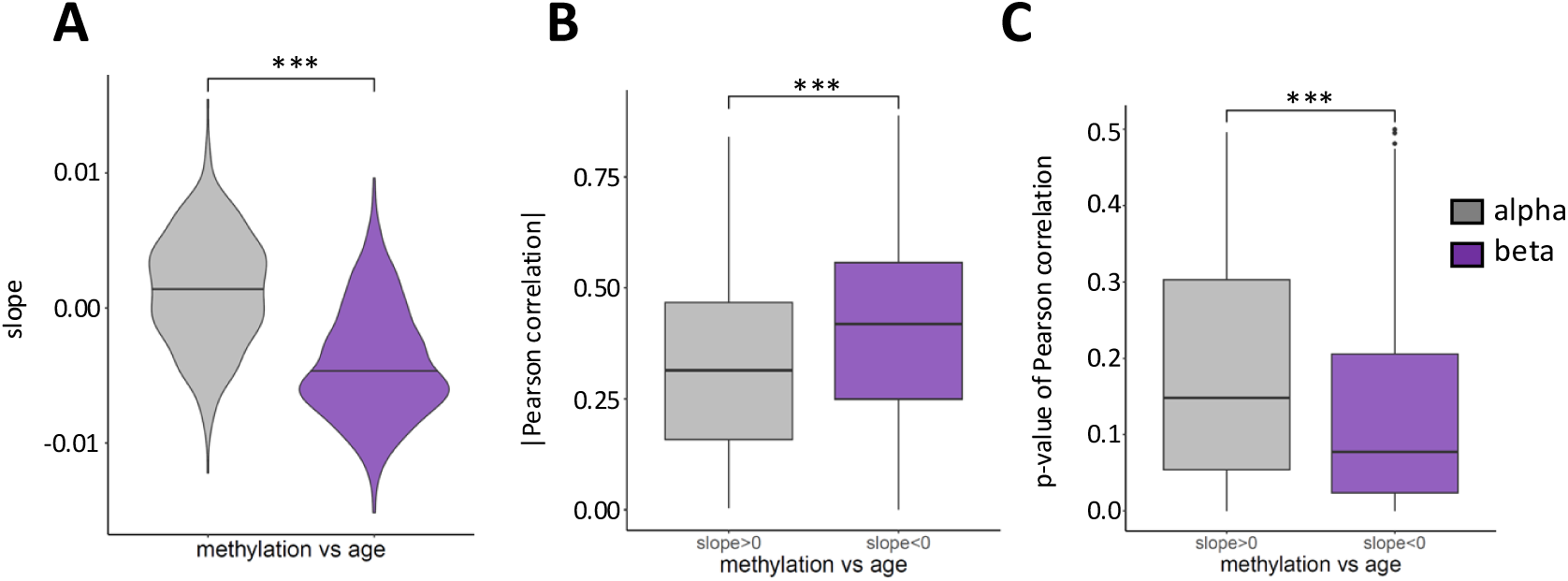
(A) Violin plots of the slopes of linear fits to methylation versus age corresponding to the regions from Fig.S5 in the two cell types. Boxplots of the absolute values of Pearson correlation coefficients (**B)** and Pearson correlation p-values (for positive association in alpha cells and negative association in beta cells) **(C)** of methylation versus age at regions with slope > 0 in alpha cells and slope < 0 in beta cells from Fig.S5. The asterisks indicate a Wilcoxon p-value < 0.001.

**Figure S7.**
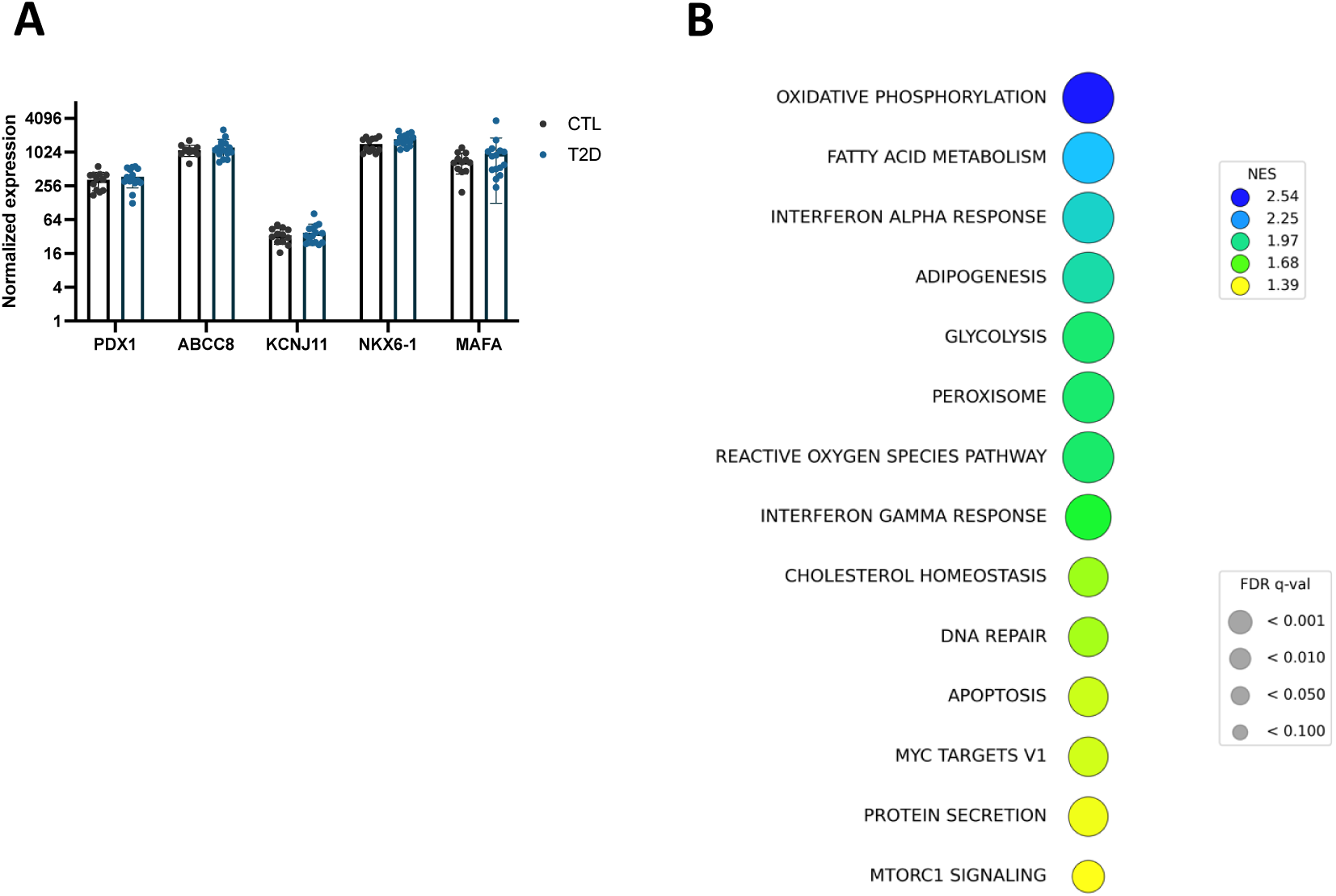
(**A**) DESeq2 normalized expression (from pseudo-bulk counts) of key beta cell identity genes in CTL (n = 13) and T2D (n = 15) donors of similar age from HPAP scRNA-Seq data, showing stable expression in T2D beta cells. (**B**) MSigDB Hallmark gene sets identified by GSEA as down-regulated in beta cells from T2D donors compared with CTLs.

**Figure S8.**
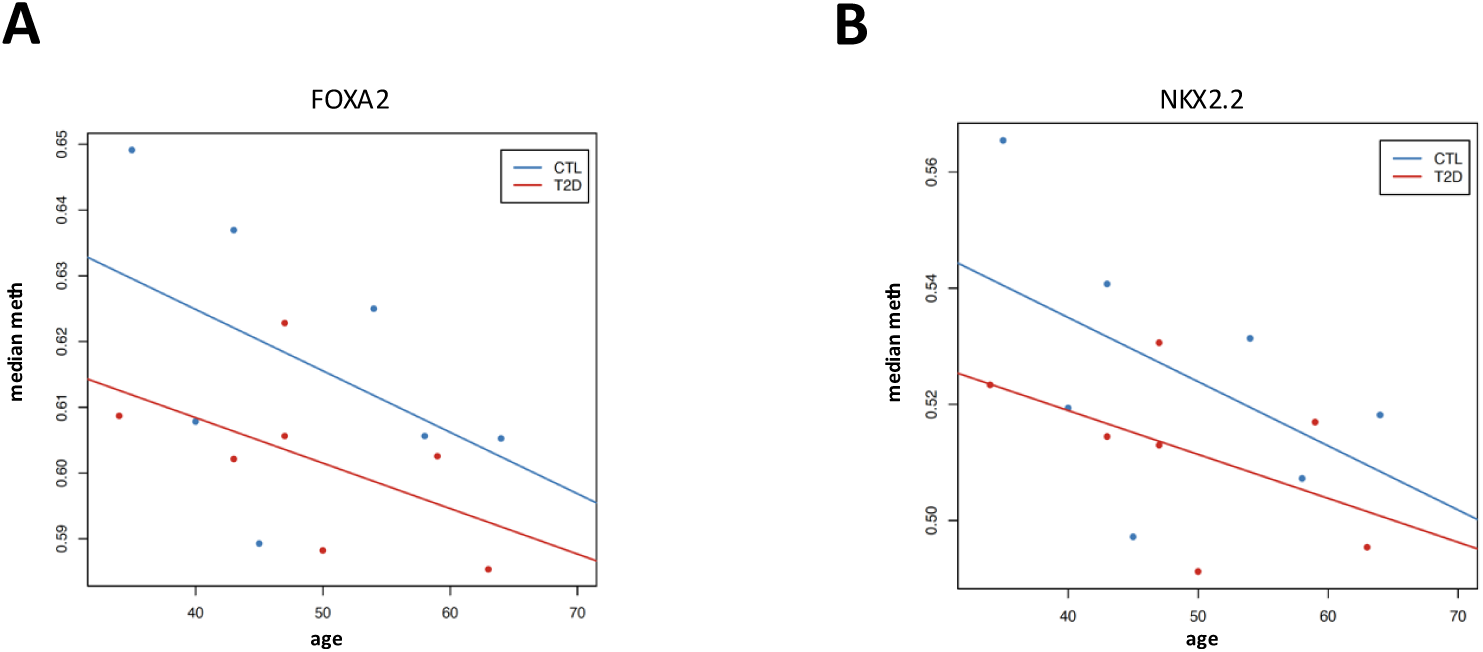
Median methylation values across the *FOXA2* **(A)** and *NKX2.2* **(B)** binding sites in the fifth quintile of the median methylation levels for 7 CTL (blue) and 7 T2D (red) age-matched donors.

## Supplemental Tables

Supplemental Table 1. Samples used in the CTL alpha vs beta cell methylome comparison.

Supplemental Table 2. Samples used in the CTL beta cell age methylome comparison

Supplemental Table 3. Samples used in the CTL alpha cell age methylome comparison

Supplemental Table 4. Samples used in the beta cell T2D vs CTL methylome comparison

Supplemental Table 5. Samples used in the alpha cell T2D vs CTL methylome comparison.

Supplemental Table 6. DMR (Differentially methylated regions) thresholds employed

Supplemental Table 7. Accession numbers for whole human islet transcription factor binding and histone modification data employed

Supplemental Table 8. Gene sets analyzed for enrichment associated with DNAm changes

Supplemental Table 9. DMR-distal genes from gene sets enriched in DMRs hypomethylated in beta cells vs. alpha cells (|delta| > 40%, FDR < 0.01)

Supplemental Table 10. DMR-distal genes from gene sets enriched in DMRs hypomethylated in alpha cells vs. beta cells (|delta| > 40%, FDR < 0.01)

Supplemental Table 11. DMR-distal genes from gene sets enriched in DMRs hypomethylated with age in beta cells (|delta| > 10%, FDR < 0.01)

Supplemental Table 12. GSEA leading edge of up-regulated gene sets with age in beta cells

Supplemental Table 13. GSEA leading edge of down-regulated gene sets with age in beta cells

Supplemental Table 14. DMR-distal genes from gene sets enriched in DMRs hypermethylated with age in alpha cells (|delta| > 10%, FDR < 0.01)

Supplemental Table 15. DMR-distal genes from gene sets enriched in DMRs hypomethylated in T2D in beta cells (|delta| > 10%, FDR < 0.05)

